# Ultrasensitive response explains the benefit of combination chemotherapy despite drug antagonism

**DOI:** 10.1101/2023.02.27.530263

**Authors:** Sarah C. Patterson, Amy E. Pomeroy, Adam C. Palmer

## Abstract

Most aggressive lymphomas are treated with combination chemotherapy, commonly as multiple cycles of concurrent drug administration. Concurrent administration is in theory optimal when combination therapies have synergistic (more than additive) drug interactions. We investigated pharmacodynamic interactions in the standard 4-drug ‘CHOP’ regimen in Peripheral T-Cell Lymphoma (PTCL) cell lines, and found that CHOP consistently exhibits antagonism and not synergy. We tested whether staggered treatment schedules could improve tumor cell kill by avoiding antagonism, using month-long *in vitro* models of concurrent or staggered treatments. Surprisingly, we observed that tumor cell kill is maximized by concurrent drug administration despite antagonistic drug-drug interactions. We propose that an ultrasensitive dose response, as described in radiology by the linear-quadratic (LQ) model, can reconcile these seemingly contradictory experimental observations. The LQ model describes the relationship between cell survival and dose, and in radiology has identified scenarios favoring hypofractionated radiation – the administration of fewer large doses rather than multiple smaller doses. Specifically, hypofractionated treatment can be favored when cells require an accumulation of DNA damage, rather than a ‘single hit’, in order to die. By adapting the LQ model to combination chemotherapy and accounting for tumor heterogeneity, we find that tumor cell kill is maximized by concurrent administration of multiple drugs, even when chemotherapies have antagonistic interactions. Thus, our study identifies a new mechanism by which combination chemotherapy can be clinically beneficial that is not reliant on positive drug-drug interactions.

## Introduction

Combination chemotherapy is essential for the treatment of many types of cancer and often consists of multiple cycles of concurrent drug administration. Several types of Non-Hodgkin Lymphoma can be cured by combination therapies built upon the four-drug regimen ‘CHOP’, consisting of cyclophosphamide (C), doxorubicin (H), vincristine (O), and prednisone (P). For example, Diffuse Large B-Cell Lymphomas (DLBCL) are commonly treated with Rituximab plus CHOP (RCHOP), and Peripheral T-Cell Lymphomas (PTCL) are commonly treated with CHOP or CHP plus brentuximab-vedotin, depending on CD30 expression^1,2^. Therapies in these CHOP-based regimens are typically administered concurrently every 14 or 21 days, for 6 to 8 cycles^3^.

Concurrent administration is in theory optimal when combination therapies have synergistic drug interactions, such that their combined effect is more than the sum of individual drugs’ efficacies. It has been recognized since the 1980s that the activity of combination therapies can depend on the timing of drug administration, with growing interest in ‘schedule-dependent synergy’ where the initial use of one drug can enhance response to another drug given some hours later^4–8^. We recently observed in DLBCL that while the RCHOP regimen is overall close to additive, the cytotoxic agents ‘CHO’ exhibit antagonistic pairwise interactions, with effects that are less than the sum of individual drugs’ efficacies^9^. Where optimizing synergy has been explored with timing adjustments on the scale of hours, drug antagonism could in principle be prevented by avoiding overlapping drug exposure, with administration separated by days. Here, we investigated the impact of drug antagonism on the efficacy of concurrent or sequential treatment by multiple drugs, using experiments and computational models. We selected PTCL as a model system because the CHOP regimen, with its several antagonistic interactions, remains the standard first-line treatment for many subtypes but cures fewer than half of patients^2^.

In this study, we confirmed that the CHOP regimen has antagonistic interactions between C-H and H-O across seven PTCL cell lines. Contrary to the expected effect of antagonism, concurrently administering CHOP achieved the most cell killing as compared to sequential administration in month-long *in vitro* experiments. These results reveal a benefit in dosing drugs together that overcomes the adverse effect of antagonistic drug interactions, which we propose can be explained by an ultrasensitive dose response model. The linear-quadratic (LQ) model, adapted from radiation oncology, describes the benefit of hypofractionation - the administration of fewer, larger doses rather than multiple smaller doses^10,11^. We adapted the LQ model to build a computational model of ultrasensitive dose response that reproduces the experimental data and shows that despite antagonistic drug interactions, concurrent therapy is more effective that sequential.

## Results

### CHOP has antagonistic interactions in PTCL cells

To test if drug antagonism observed in DLCBL also occurs in PTCL, we measured pharmacodynamic drug interactions within the CHOP regimen in seven PTCL cell lines, representing four histologically diverse subtypes (Anaplastic Large Cell Lymphoma (ALCL), Natural Killer (NK), T-cell Large Granular Lymphocytic (T-LGL), and PTCL-not otherwise specified (PTCL-NOS)). Pairwise drug interactions were measured in microtiter plates with 11×11 drug-drug concentration gradients spanning clinically relevant concentrations by reference to C_sustained_, the observed concentrations in patient serum 6 hours after administration^12^. High-order combinations (three or four drugs) were measured as a dose-response to a mixture with fixed concentration ratio based on C_sustained_. Relative cell viability was quantified by an ATP-based luminescence assay, which provided a 10,000-fold dynamic range in PTCL cell lines (100% to 0.01% relative live cell count) (**Supplemental Figure 1**). Cyclophosphamide is a pro-drug and was substituted with 4-hydroperoxy-cyclophosphamide, which generates the active metabolite 4-hydroxy-cyclophosphamide in buffered media ^13^. Pairwise interactions with prednisolone (the active metabolite of the prednisone) were not studied because the drug lacked *in vitro* activity in all PTCL cell lines, as reported for DLBCL cell lines^14,15^. Drug interactions were analyzed by two common metrics: Loewe’s dose additivity model (**Figure 1**) and Bliss’ independence model (**Figure 2**)^16^. Isobologram analysis, based on Loewe’s method, provided a detailed view of pairwise interactions, whereas the Bliss model readily extends to high-order interactions and was used to analyze combinations of two, three, and four drugs. To overcome limitations of these single drug interaction metrics, we also applied the ‘Multidimensional Synergy of Combinations’ (MuSyC) framework which unifies the methods of Loewe and Bliss (**Supplemental Figure 2**)^17^.

**Figure 1.**
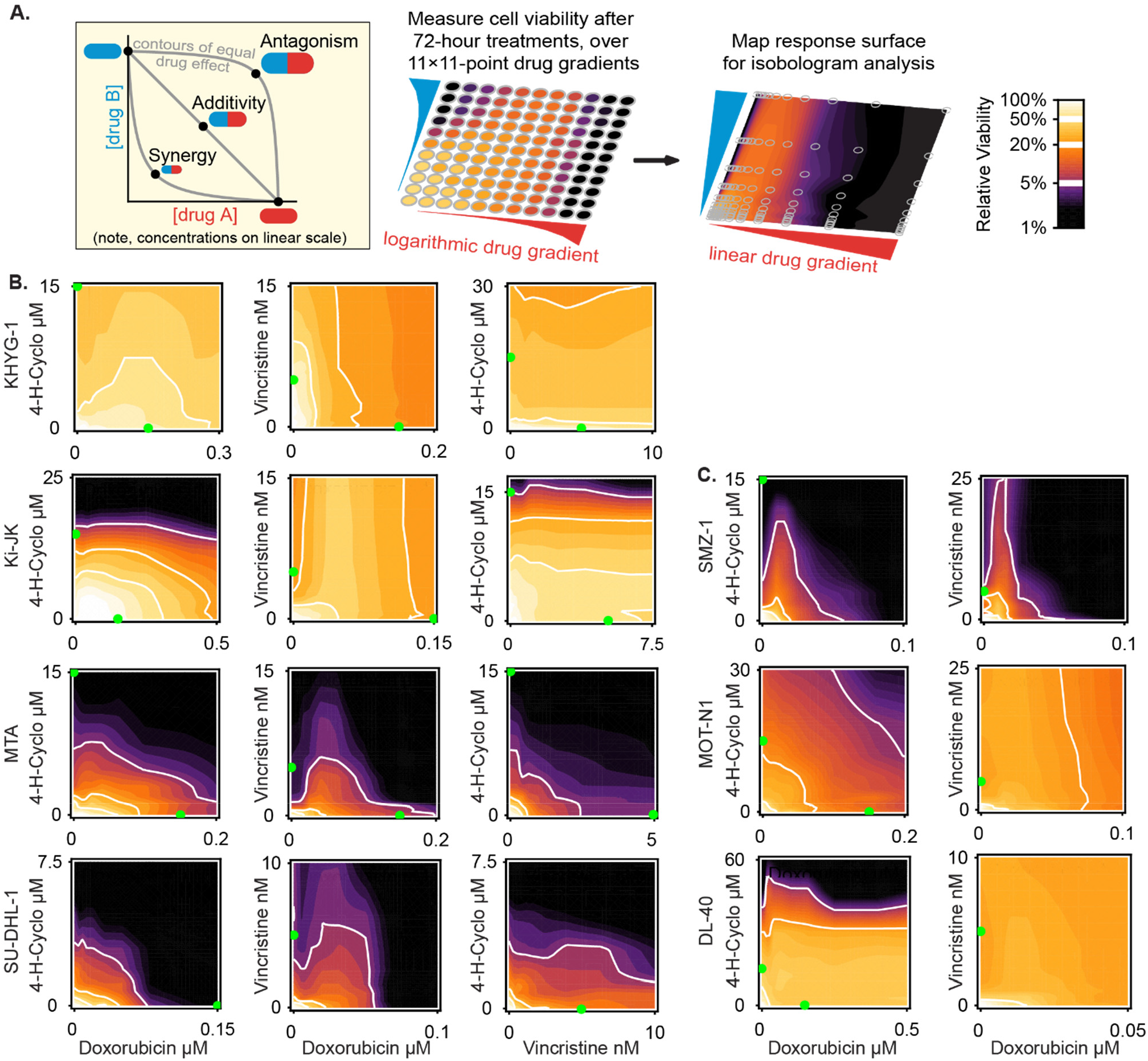
Isobologram analysis shows antagonism between CH and HO in the CHOP regimen for Peripheral T-Cell Lymphoma. A) Left: shapes of equal-effect contour lines (isoboles) reveal non-additive drug interactions (Palmer et al.)^9^. Right: Cell viability is measured across an 11×11 concentration gradient to graph isobolograms. B) CH, HO, and CO drug interactions in four Peripheral T-Cell Lymphoma (PTCL) cell lines: KHYG-1, Ki-JK, MTA, SU-DHL-1 (n=4 replicates per plot). White contour lines highlight 50%, 20%, and 5% relative viability. Green dots mark a clinically relevant concentration, C_sustained_ (the average plasma concentration in humans 6 hours after administration^12^). C) CH and HO pairwise drug interactions in three additional PTCL cell lines: SMZ-1, MOT-N1, DL-40 (n=4 replicates per plot).

**Figure 2.**
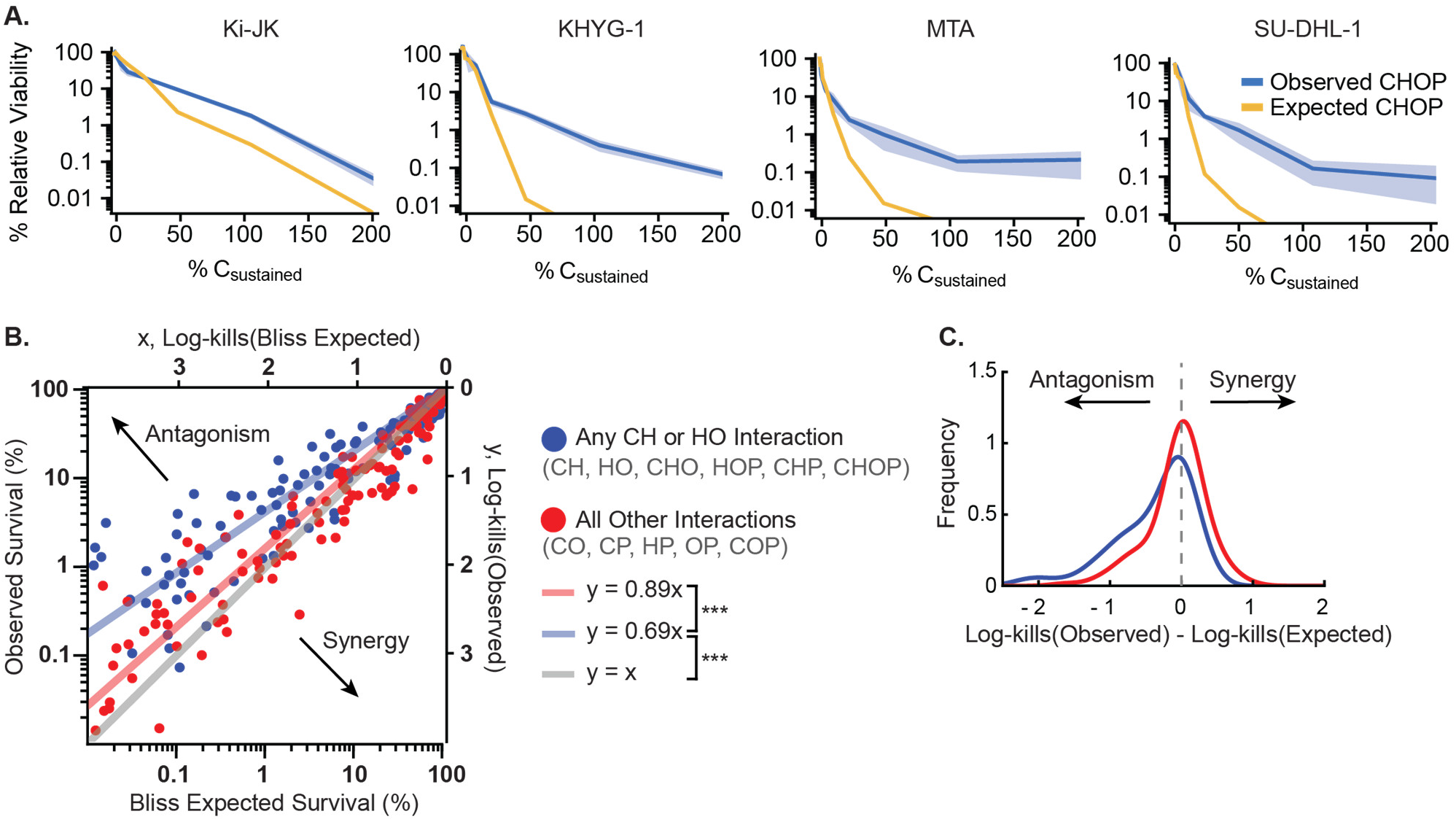
Analysis of drug pairs, triplets, and quadruplet CHOP confirms antagonistic interactions by the Bliss Independence model. A) Effect of CHOP combination measured at concentrations from 0 to 200% of C_sustained_, compared with Bliss Independence model computed from monotherapy effects, in four PTCL cell lines: KHYG-1, Ki-JK, MTA, SU-DHL-1. Blue shading represents 95% confidence interval (n=8 replicates). B) Relationship between observed cell viability, and predicted viability from Bliss Independence model, for all possible pairs, triplets, and quadruplets of drugs in CHOP across four PTCL cell lines (n=237 points across concentration range up to 500% C_sustained_; each point is an average of 8 measurements). All combinations containing antagonistic pairs CH or HO are colored in blue; all other combinations in red. Lines of best fit show that, as a group, combinations containing the drug pairs CH or HO (blue line) are antagonistic compared with the Bliss independence model (black line) (*P* = 6×10^−34^, Student’s t-test, n=117), and also more antagonistic than all combinations without CH or HO (red line) (*P* = 3×10^−14^, Student’s t-test, n=237). Overall, all CHOP measurements (blue and red combined) are antagonistic compared with the Bliss independence model (black line) (*P* = 7×10^−35^, Student’s t-test, n=237). C) Histogram depicting the difference in observed and expected log-kills (-log_10_(relative viability) for all interactions involving CH or HO (blue), and all remaining interactions (red)).

Loewe’s dose additivity model assesses potency of response (such as IC50) and is depicted by ‘isobologram plots’ which show contours of equal effect (isoboles) (**Figure 1a**)^18,19^. If drugs A and B are additive, the isoboles will be straight lines as half of the IC50 of drug A plus half of the IC50 of drug B constitutes the IC50 of the combination. A drug combination is synergistic if potencies are enhanced in combination (convex isoboles) and antagonistic if potencies are diminished in combination (concave isoboles). Isobologram analysis confirmed that CHOP exhibits antagonistic interactions in PTCL cell lines (**Figure 1b**). Specifically, the activity of vincristine was suppressed by doxorubicin in six of seven cell lines, with the exception being a cell line in which vincristine lacked activity. Cyclophosphamide and doxorubicin (C-H) were additive in two cell lines, less than additive in two cell lines, and suppressive (weaker than monotherapy) in two; one cell line had little response to doxorubicin so the interaction was not assessable. Cyclophosphamide and vincristine (C-O) were approximately additive when active in four cell lines and not studied further. No synergistic interactions were observed in any cell line.

The Bliss independence model, which assesses magnitude of effect, also classified the CHOP regimen as less than additive (**Figure 2**). In the Bliss model drugs are non-interacting if their combined effect is consistent with independent probabilities of cell death. For example, if 10% of cells survive Drug A and 10% of cells survive Drug B, the Bliss model predicts that 1% of cells will survive the combination A+B (10% x 10% = 1%); this corresponds to addition of ‘log-kills’ (90% inhibition = 1 log-kill)^20^. Drug combinations are classified as synergistic or antagonistic if their effect is greater or lesser than expected. At clinically relevant concentrations, the full CHOP combination was less than additive in each of four PTCL cell lines, with up to 100-fold more lymphoma cell viability than expected (**Figure 2a**). Dose-responses to every pair, triplet, or quadruplet of drugs in CHOP (11 combinations) were also measured in four cell lines and compared to the Bliss model (**Figure 2b**). Globally, the relation between observed and expected cell viabilities was described by linear regression with a slope of 0.79, which is significantly different from 1.0 as would be expected by Bliss independence (*P* = 7×10^−35^). This slope indicates that combinations generally produced ~20% fewer log-kills than expected. Combinations containing H-O and C-H were the most antagonistic, with a slope of 0.69 indicating ~30% fewer log-kills than expected (*P* = 6×10^−34^), and a significantly stronger degree of antagonism than other subsets of drugs in CHOP (**Figure 2c;** *P* = 3×10^−14^).

Finally, drug interactions were assessed by the MuSyC model which quantifies changes in both potency and efficacy, to unify Loewe’s and Bliss’ models^17^. MuSyC analyzes interactions among drug pairs by fitting to sigmoidal dose-response surfaces; we extended the method to analyze higher-order combinations without requiring sigmoidal fitting (**Methods**). Briefly, we calculated the Bliss expected combination response from monotherapy data, and then applied changes in potency and magnitude of effect to best fit the experimentally measured dose response. This defines changes in potency and efficacy compared to the expected additive response (**Supplemental Figure 2**). All drug combinations except one were additive or antagonistic, with up to two-fold decreases in potency and/or efficacy in combinations containing CH and HO. The sole exception was vincristine and prednisolone (O-P) in the Ki-JK cell line, in which prednisolone lacked individual activity but enhanced vincristine response. Overall, three forms of analysis consistently found that drugs in CHOP are predominantly additive or less when combined on PTCL cells, with notable antagonism between C-H and H-O.

### Concurrent therapy is more effective than sequential

All metrics of drug interaction confirmed that antagonism between H-O and C-H decreases the overall efficacy of CHOP. We hypothesized that separating the administration of antagonistic drugs would increase the efficacy of the combination *in vitro*. We compared the long-term efficacy of concurrent or sequential therapy by measuring live cell count throughout two 12-day treatment cycles (**Figure 3a**). In concurrent therapy, all four drugs in CHOP were administered at the start of each cycle; this resembles the clinical regimen. In sequential therapy, three drugs were administered at the start of each cycle and the remaining drug was given mid-cycle. Total administered dose was constant across all conditions. MTA cells were studied because their levels of drug sensitivity and drug interactions were typical of these PTCL cell lines. Concurrent therapy was observed to produce the greatest therapeutic effect, whereas sequential regimens that offset C or H were much less effective, leaving 10 times more surviving lymphoma cells compared to concurrent therapy (**Figure 3b**). Thus, antagonistic drug interactions did not compromise the activity of concurrent therapy.

**Figure 3.**
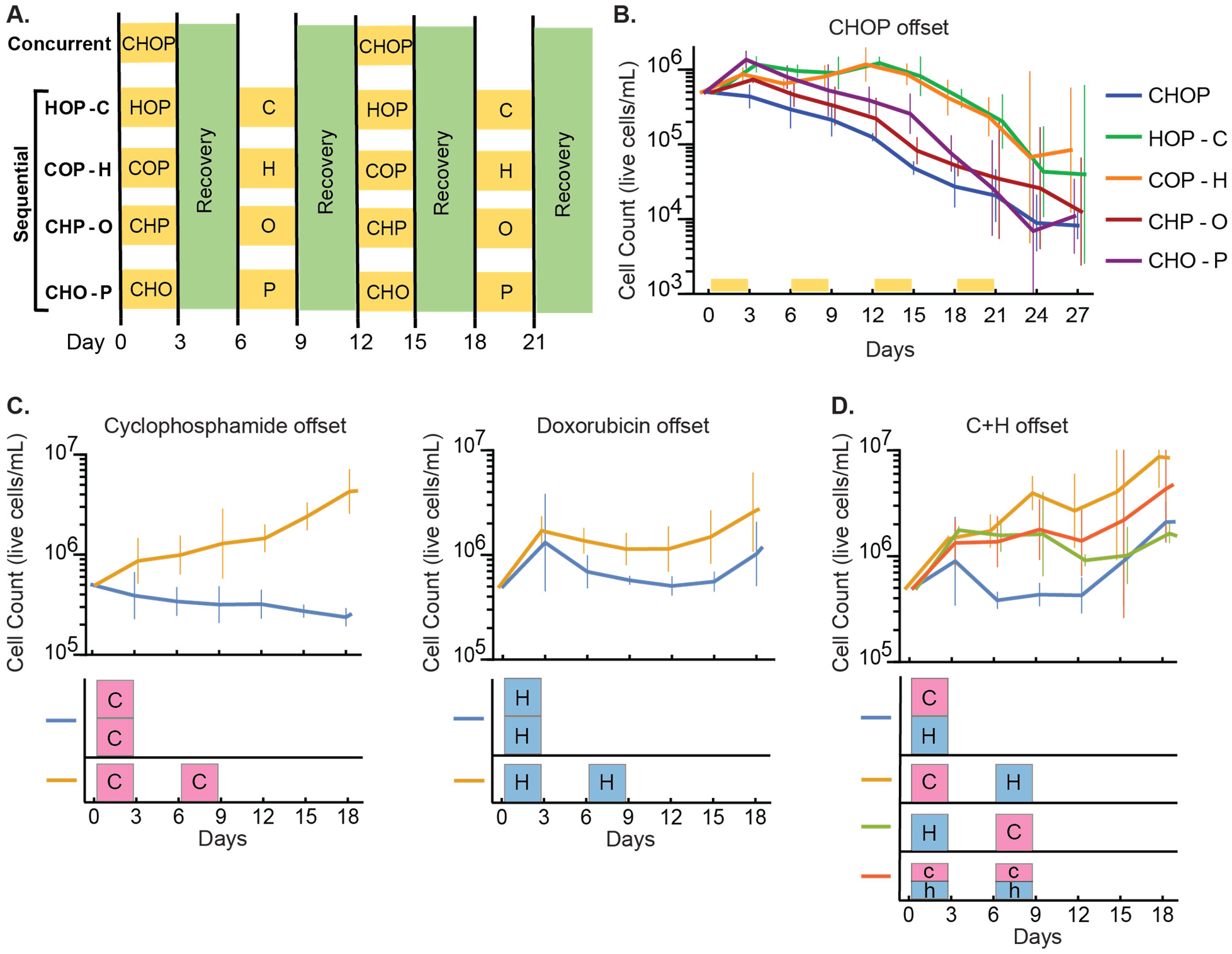
Concurrent administration of CHOP is more effective in vitro than sequential administration. A) Experimental design for concurrent versus sequential treatment schedules. PTCL cultures (MTA) were treated in 72-hour blocks, alternating between treatment, and recovery in drug-free media. Drugs were administered at an equipotent ratio (**Methods**). B) Live cell count (hemocytometer and trypan blue) over the span of two cycles of concurrent or sequential CHOP. Yellow bars indicate drug treatment periods (n=3 independent cultures per condition, error bars are 95% confidence). C) Live cell count after treatment with two simultaneous doses or two sequential doses, of 4-H-cyclophosphamide only (1 dose = 2 μM), or of doxorubicin only (1 dose = 20 nM). Treatment schedules are visualized underneath each time axis (letters C and H represent one dose). D) Live cell count after CH combination treatment using concurrent, sequential, or ‘split’ dosing (both drugs at half dose, for two sequential doses). Lowercase c and h in the schematic represent half-doses: c = 1 μM 4-H-cyclophosphamide; h = 10 nM doxorubicin.

We hypothesized that the dose intensity of concurrent therapy may confer an advantage that overcomes antagonistic drug interactions. First, to test this hypothesis without the confounding effect of drug interactions, we compared concurrent and sequential treatments of single agent cyclophosphamide (C) or doxorubicin (H), the drugs that had the most pronounced difference when administered sequentially. Concurrent therapy consisted of two doses at the beginning of the cycle, and sequential therapy consisted of one dose at the beginning of the cycle and one mid-cycle (**Figure 3c**). Even as monotherapies, administering the entire dose at one time had a 5- to 10-fold greater effect than dividing the same total dosage between two treatment times. This data shows a benefit to high intensity dosing that is independent of drug interaction. Second, to test whether this advantage applies even to an antagonistic drug combination, we compared concurrent therapy with C and H to sequential treatments of C then H or H then C, separated by three days. Additionally, a split dosing regimen was evaluated consisting of C plus H each administered at half concentration (**Figure 3d**). By day 12 (6 days after the second dose), both sequential treatments and the split treatment were significantly less effective than concurrent administration of C and H. Thus, for both monotherapy and combination therapy, concurrent administration of the greatest possible dose intensity produces the greatest inhibitory effect, despite antagonistic drug interactions.

An inherent benefit of concurrent drug administration could be explained by two hypotheses. First, tumor cells could have physiological responses to the first dose of chemotherapy that make them more resistant to future treatments, a form of adaptive resistance that would cause sequential therapy to be inferior. Second, the dose response function could exhibit a form of ultrasensitivity where higher doses produce a disproportionately large increase in cytotoxic effect, thus favoring concurrent over sequential therapy. Recognizing these are not mutually exclusive, we next tested whether either the adaptive resistance hypothesis or ultrasensitive response hypothesis can explain the benefit of concurrent therapy.

### Adaptive resistance does not cause the deficiency of sequential therapy

An adaptive cellular response to an initial chemotherapy dose may diminish sensitivity to subsequent treatments (as distinct from long-term evolutionary selection). We tested this hypothesis by treating MTA cells with partially inhibitory doses of single agents C, H, O, or placebo, for three days, and then after four days of recovery, repeated dose response measurements. In no case were pretreated cultures less sensitive to subsequent therapy (**Supplementary Figure 3**). Therefore, shortterm adaptive resistance does not explain the superiority of concurrent CHOP administration.

### Ultrasensitive response can explain the advantage of concurrent therapy

Like many chemotherapies, the agents in CHOP exhibit approximately exponential dose response functions over the first ~90% of inhibition^21^. This corresponds to a linear increase in log-kills with drug concentration (**Supplementary Figure 4**), which would predict equal effect of concurrent or sequential therapy for the same total dosage. However, if this relationship is ultrasensitive (more than linear) at greater cytotoxic effect, then the higher dose-intensity of concurrent therapy could be advantageous. Ultrasensitive dose responses are a central concept in radiation oncology, where the Linear-Quadratic (LQ) model describes two components of dose response: a linear term describing the probability of cell death per single event of DNA damage, and a quadratic term describing the added probability of death from an accumulation of DNA damage (**Methods, Equation 1**). The LQ model describes the relative efficacy of hyperfractionated radiation (many small doses) versus hypofractionated radiation (one or few large doses) in various scenarios. When the quadratic term is large, the greatest cytotoxicity is achieved by a single high dose of radiation, akin to concurrent administration of many drugs. Here we apply the LQ model to cytotoxic chemotherapies that also target DNA, to test the hypothesis that ultrasensitive dose response explains the benefit of concurrent chemotherapy.

In a linear dose response, doubling the dose that produces one log-kill will produce two log-kills (**Figure 4a**). In principle the same effect would result from sequential administration of two such doses, because 1 log-kill followed by 1 log-kill is a net effect of 2 log-kills. Conversely, with an ultrasensitive response as described by the LQ model, concurrent administration (doubling the total dose) could produce as much as 4 log-kills, providing ‘increasing returns’ (**Figure 4a**). However, when examined on a log-scale, dose responses to single or combined agents in CHOP were sublinear in each of 7 cell lines, i.e., exhibiting decreasing returns (**Figure 4b and Supplemental Figure 4**). In a sub-linear response, doubling the dose that produces one log-kill produces less than two log-kills; this would favor sequential therapy, in contradiction to the experimentally demonstrated superiority of concurrent therapy.

**Figure 4.**
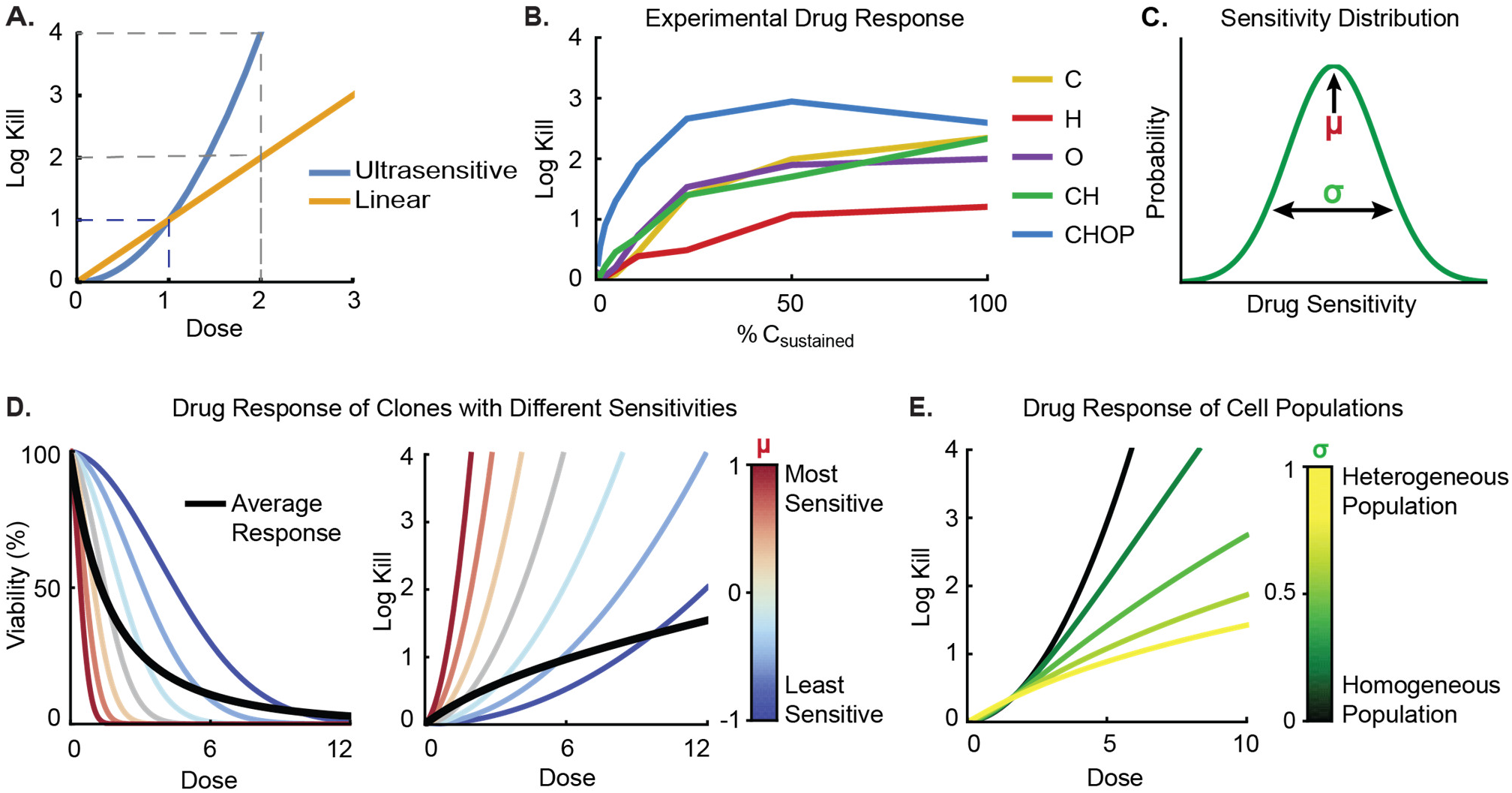
Ultrasensitive dose responses will be masked by cellular heterogeneity. A) A comparison of linear and ultrasensitive dose response functions. With a linear response, doubling the concentration for 90% inhibition (1 log-kill) achieves 99% inhibition (2 log-kills). With an ultrasensitive response, doubling the dose more than doubles the number of log-kills. B) CHOP and its components exhibit ‘sub-linear’ dose responses in PTCL cell lines (n=8) (MTA shown here; see other cell lines in **Supplemental Figure 4**). C) Phenotypic heterogeneity, which can be genetic or stochastic in origin, can cause drug sensitivity to vary between individual cells. Variance in sensitivity to drugs in CHOP is observed to follow a log-normal distribution which is quantified by the mean (μ) and standard deviation (σ) of drug sensitivity (**Supplemental Figure 5**). D) Ultrasensitive dose responses with variable sensitivity illustrated by seven example clones 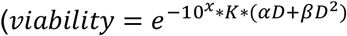 see **Methods**). Drug sensitivity is indicated by color where red represents more sensitive, and blue represents less sensitive. The black line is the dose response of a whole population, composed of variable clones with log-normally distributed drug sensitivities. Note that the population’s average response has a different shape from individual clones, when graphed on a log-scale. This arises because the average of 1 log-kill and 3 log-kills is 1.3 log-kills (average of 10% and 0.1% is ~5%). E) Increasing cellular heterogeneity causes ultrasensitive dose responses to be obscured at the population scale. Here, all cells have ultrasensitive dose responses, but this is only evident in homogeneous populations (dark green line). Heterogeneous populations show sub-linear dose responses despite underlying ultrasensitivity of single cells (yellow line).

To resolve this seeming paradox we turned to research on cell-to-cell variability, which has shown that population-level dose responses are an average of single-cell phenotypes^22^. We hypothesized that individual cells could have ultrasensitive dose responses that appear more gradual at the population level due to cellular heterogeneity. Indeed, we recently observed variability in the sensitivities of lymphoma cells to the drugs in CHOP, with a log-normal distribution of sensitivities revealed by high-complexity clone tracing as well as CRISPR screens (**Supplemental Figure 5**)^9^. We therefore built a mathematical model of ultrasensitive dose response in a population of cells having log-normally distributed drug sensitivity (**Figure 4c, Methods**). In this model, cells that are more or less drug sensitive have the same dose response function with the sole difference of shifting left or right (respectively) along a concentration axis (**Figure 4d**). This model showed that when individual cells have ultrasensitive dose responses (increasing returns) but variable drug sensitivity, the population-level response can show diminishing returns (**Figure 4e**). An underlying ultrasensitivity would only be apparent in a homogenous population of cells. In this theory, the appearance of ‘diminishing returns’ is not because chemotherapy efficacy declines with higher dosage, but because the remaining few cells are especially hard to kill. We next tested whether this theory can reconcile three seemingly contradictory observations: that antagonistic chemotherapies, showing diminishing returns in their dose response, could be maximally effective when administered concurrently.

Given two model features – cellular heterogeneity and ultrasensitive dose response – we tested whether either or both could explain observed dose responses and dynamics of sequential or concurrent therapy. We simulated cell population dynamics in response to treatments that induce a fraction of cell death, according to dose response functions which include antagonistic interaction between C and H. All model parameters were measured except for the degree of ultrasensitivity (β) and variance in drug sensitivity (σ), which were fitted to experimental data (**Methods and Supplemental Figure 6**).

A ‘null model’ with neither heterogeneity nor ultrasensitivity failed to reproduce any of the observed data (**Figure 5**). A model including only cellular heterogeneity only reproduced the diminishing returns of dose response, and a model including only ultrasensitivity only reproduced the advantage of concurrent therapy. Finally, a model including both heterogeneity and ultrasensitivity was able to reproduce both the population-level dose response as well as the observed advantage of concurrent therapy over sequential therapy, even with the model’s inclusion of antagonistic drug interaction (**Figure 5**).

**Figure 5.**
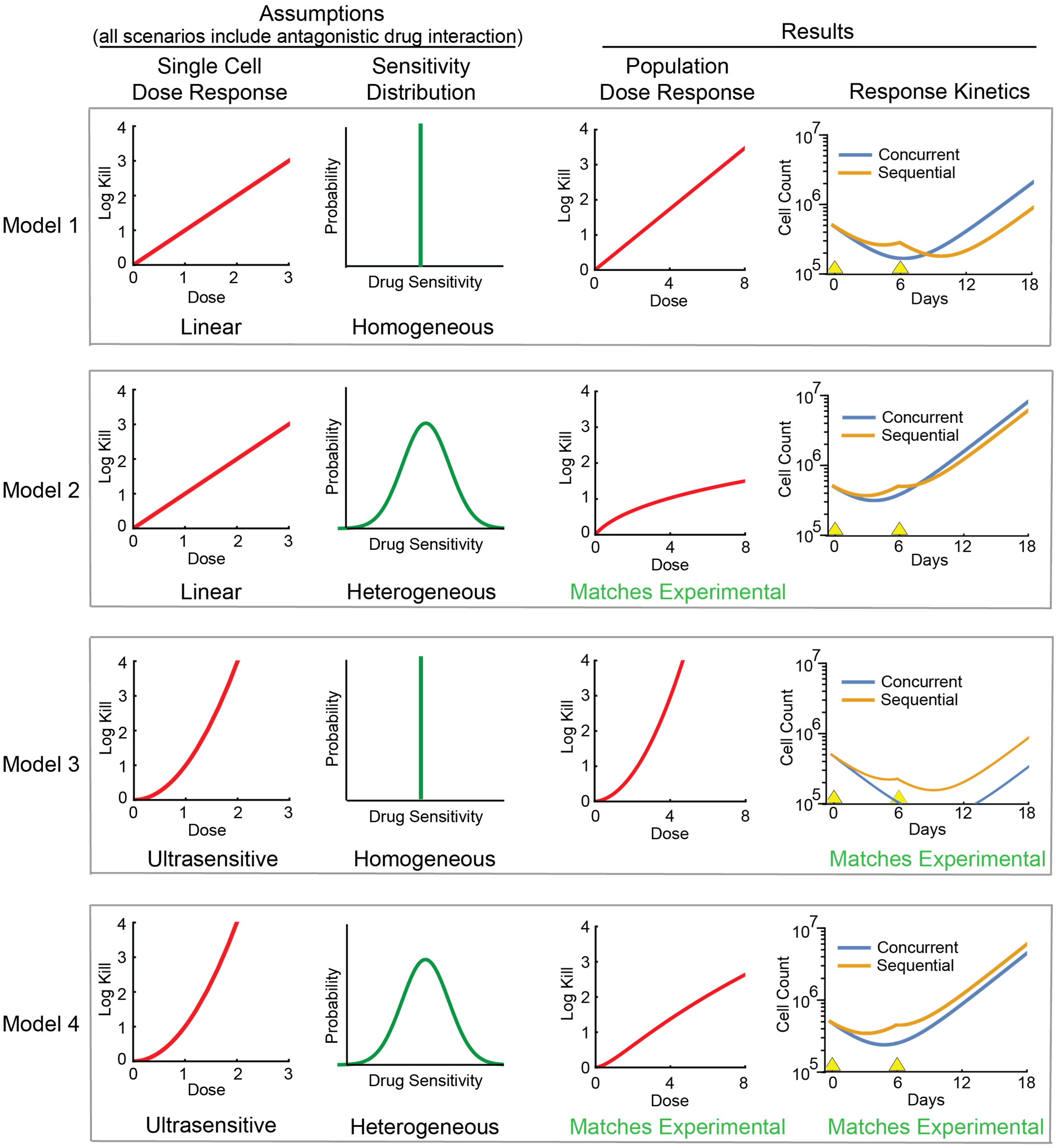
Ultrasensitivity and cellular heterogeneity can together explain the observed advantage of concurrent combination chemotherapy. Four mathematical models of tumor response to concurrent or sequential chemotherapy are compared, based on linear or ultrasensitive dose responses, and homogeneous or heterogeneous cell populations. Models were categorically judged by their ability to reproduce the observed population-level dose response (**Figure 4b**), and dynamics of response to concurrent or sequential use of the antagonistic CH drug pair (**Figure 3d**). All models include the observed antagonism between C and H (see **Methods** and **Supplemental Figure 6** for parameters). Yellow triangles mark treatment times (concurrent: C and H both on day 0; sequential: C on day 0 and H on day 6).

### The effect of drug interactions on optimal schedule

We next used this model to simulate how various drug-drug interactions affect concurrent versus sequential treatments. For drug interactions ranging from synergistic to antagonistic, we compared responses to drug pairs where the same total doses are applied but with different schedules: concurrent, sequential, or split (half doses given twice as often). In scenarios implementing synergy or additivity, the greatest tumor cell kill was achieved by the high dose intensity of concurrent therapy (**Figure 6a**). This simulated effect of additivity (which also applies to a drug ‘combined’ with itself)^23^ resembles the experimentally observed responses to single agents 4-H-cyclophosphamide or doxorubicin at full dose versus sequential half doses (**Figure 3c**). In the presence of mild antagonism, concurrent administration is superior but to a lesser degree, as sequential therapy has some benefit from avoiding antagonism (**Figure 6a**). This model resembles experimental observations for the mildly antagonistic combination of 4-H-cyclophosphamide and doxorubicin (**Figure 3d**). With sufficiently strong antagonism, a point is reached where avoiding antagonism matches the benefit of high dose intensity, making sequential and concurrent schedules similarly effective (**Figure 6a**). This consequence of strong antagonism corresponds to the experimental observation that concurrent CHOP was similar in efficacy to the sequential regimen ‘CHP then O’, which avoided the strong antagonism of vincristine by doxorubicin (**Figures 1, 3b**). Therefore, when drug antagonism is present in combination regimens, its detrimental effect can be counterbalanced by the benefit of higher dose intensity, and conversely, the benefit of avoiding antagonism by sequential regimens may be balanced by the loss of dose intensity.

**Figure 6.**
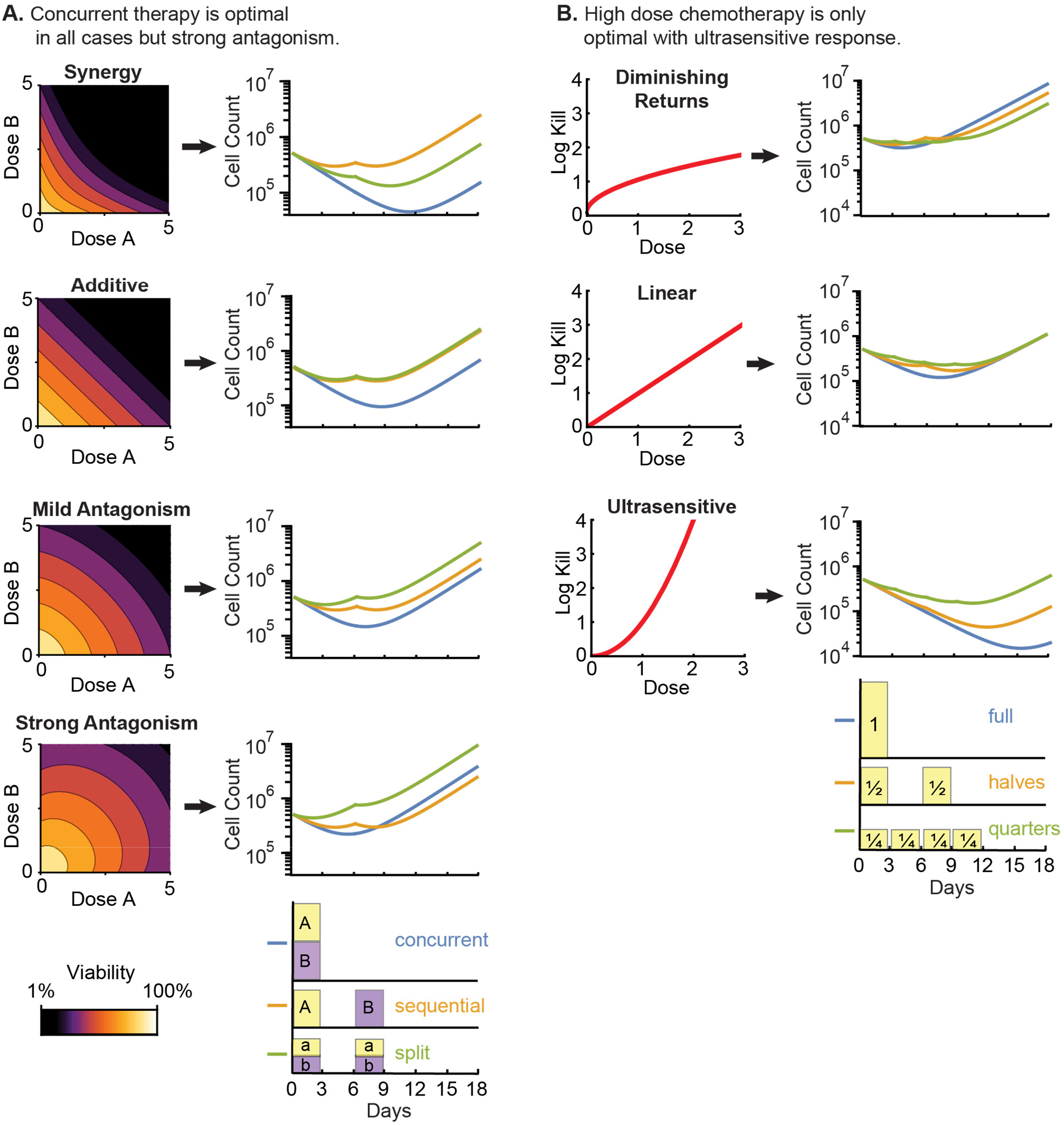
Effect of drug interactions and dose intensity on sequential and concurrent chemotherapy. A) Effect of drug interactions on efficacy of different treatment schedules. Simulated dynamics of response to different treatment schedules for a drug pair ‘A+B’ exhibiting synergy (i = 1.5), additivity (i = 0), mild antagonism (i = −0.8), and strong antagonism (i = −1.5). Isobolograms are shown on the left, and live cell dynamics are shown on the right, for concurrent use of both drugs (blue), sequential use of drugs (orange), or split dosing (both drugs at half dose, two sequential doses; green). B) Effect of dose response shape on efficacy of different treatment schedules. Here we consider only monotherapy to remove influence of drug interactions. Live cell dynamics are simulated for dose response functions that either produce diminishing returns 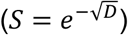; linear response (*S* = *e^−D^*), or ultrasensitive response 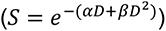. Dose response functions are shown on the left, and live cell dynamics are shown on the right for intense treatment, consisting of the full dose on day 0 (blue), sequential use of two half-doses (orange), or sequential use of four quarter-doses (green). All models in both panels feature a consistent distribution of drug sensitivity (**Methods**).

### High-intensity chemotherapy is only optimal with ultrasensitive dose response

Finally, we investigated how the shape of dose response affects the relative efficacy of high-intensity versus low-intensity chemotherapy schedules. We simulated single-agent treatments (avoiding drug interaction) where the same total dosage is either concentrated or spread over time. Specifically, we test administering therapy all at once (day 0), as two half doses (days 0 and 6), or as four quarter doses (days 0, 3, 6, 9). When the dose-response function is sub-linear (diminishing returns), the prolonged low-dose regimen is the most effective –opposite to experimental results (**Figures 3c, 6b**). With a linear dose response, efficacy depends only on total dose and all schedules have an identical final result, which is also inconsistent with experiments. Only with a supra-linear or ultrasensitive dose response is the high-intensity schedule most effective, with prolonged low-dose chemotherapy being far inferior. This is the only scenario that is consistent with the experimentally observed inferiority of lower-intensity chemotherapy (**Figures 3c, 6b**). Collectively these results suggest that cellular dose responses must be ultrasensitive in scenarios where high-intensity chemotherapy is superior to prolonged low-intensity chemotherapy, such as curative treatments for aggressive lymphomas.

## Discussion

Across a variety of T-cell lymphoma cell lines, the drugs comprising the clinically standard CHOP regimen exhibited antagonistic interactions, which diminish the efficacy of combination therapy. However, despite these antagonistic drug interactions, we observed that concurrent administration of all agents in CHOP produced superior or equal efficacy compared with sequential regimens. We propose that these surprising observations can be explained by an ultrasensitive dose response to chemotherapies, such that maximal efficacy results from the highest intensity regimen, whereas inferior efficacy is produced when the same total dosage is spread out over time in sequential or ‘hyperfractionated’ regimens. These findings suggest that ultrasensitive dose response is a mechanism contributing to the benefit of combination chemotherapy, distinct from synergistic drug interaction, that can overcome the detrimental effect of antagonistic drug interactions. This phenomenon may contribute to the observed clinical superiority of cycles of high-dose chemotherapy in multiple types of cancer, in contrast to the relatively scarce use of prolonged low-dose chemotherapy^24–28^. Ultrasensitive dose responses may also provide a mechanistic explanation for clinical trials in which concurrent regimens have outperformed sequential regimens^29–32^, although these results are also affected by tolerability.

Ultrasensitive dose responses are expected if cells become more likely to die as chemotherapy-induced damage accumulates; the same phenomenon is recognized in radiotherapy by the LQ model. We also show that heterogeneity in drug sensitivity will cause *populations* of cells to exhibit linear or sub-linear dose responses even when *individual* cells possess ultrasensitive (supra-linear) responses. This arises because some drug sensitive cells are easily killed, and remaining resistant cells are difficult to kill, such that the cytotoxic effect in a heterogeneous population shows diminishing returns with increasing dose. Indeed, the chemotherapies in CHOP exhibited sub-linear dose responses in T-cell lymphoma cultures. This challenge cannot be resolved by single-cell measurements, as dose-response functions cannot be measured in the same single cell – once a cell dies it cannot be exposed to other doses. Even a population derived from a single clone will not consist of cells with identical drug sensitivity, as phenotypic heterogeneity can arise rapidly due to stochastic fluctuations. Given the fundamental inability to directly observe this phenomenon, we instead show by simulation that *without* ultrasensitive dose responses, prolonged low-dose chemotherapy would be equal or superior to high-dose chemotherapy. This suggests that in clinical scenarios where cycles of intensive chemotherapy are more effective than prolonged low-dose therapy, such as aggressive lymphomas, cancer cells logically must have ultrasensitive dose response functions. These results align with the existing theory that a drug’s dose response function can inform its optimal treatment schedule.^33^

The insights from CHOP may apply to other cytotoxic chemotherapies, which are often administered as cycles of high doses, and often directly or indirectly damage DNA akin to the DNA-damaging effects of radiation in the LQ model. Conversely, we anticipate that ultrasensitive responses are unlikely to apply to treatments that are dosed daily to achieve sustained inhibition of oncogenic signaling, such as kinase inhibitors and hormone therapies. The character of dose responses may be relevant to designing novel combination regimens, since when ultrasensitivity is present, tolerable combinations of active therapies have promise even without positive drug-drug interactions.

Tolerability is an overriding consideration in cancer treatments whose importance is emphasized by our findings. Ultrasensitive dose responses suggest that although combinations of active therapies may increase efficacy when tolerable, these advantages may be seriously compromised when toxicity necessitates dose reductions. In particular, loss of efficacy may be non-linear with respect to reduction in dose. Aggressive lymphomas have a history of more intensive chemotherapy regimens failing to improve survival, possibly because more drugs did not correspond to a higher achievable sum of dose intensities^34^. In PTCL, the addition of romidepsin to CHOP necessitated more frequent dose reductions of the CHOP backbone, which potentially explains the lack of survival improvement^35,36^. Conversely, successful combinations such as Rituximab plus CHOP for DLBCL have been well tolerated. In short, when high-dose intensity is important, tolerability is important.

The chief limitation of this study is that the dose response function of any given single cell is fundamentally unobservable, and therefore the evidence for ultrasensitive dose responses comes from their consequences for high versus low-intensity, and concurrent versus sequential regimens. The most compelling evidence for this theory is the finding that prolonged low-dose chemotherapy would hypothetically be the optimal use of chemotherapy if the dose responses of single cancer cells were sub-linear. Our study is limited to the cytotoxic agents in CHOP, because prednisolone (the active metabolite of prednisone) has little to no single agent activity in PTCL cell lines, as also reported for DLBCL^9,15^. Finally, we have not investigated the mechanistic causes of antagonistic drug interactions within CHOP, as the purpose of this study was to understand the consequence of observed drug interactions on treatment schedules. Similarly, some temporal effects of drug interactions are not understood, such as our observation that H before C was more effective than C before H (**Figure 3d**). Since concurrent therapy was the most effective use of CHOP, these remaining questions do not have clear importance.

Models to optimize treatment schedules have an established history in radiation oncology, and may also be useful for chemotherapies as discussed by McKenna et al^37^. The utility of cycles of high-dose chemotherapy, compared for example with daily low-dose chemotherapy, is well recognized and established in oncology. Our study found that this advantage is not obvious when considered quantitatively, especially in light of antagonistic drug interactions, but can be rationalized by heterogeneity in dose responses. This work adds to our understanding of the therapeutic efficacy of existing treatments, which can form a basis for future prospective applications. The framework presented here can be adapted to other therapies and cancer types, to understand the effects of dose intensity, treatment schedules, and drug interactions.

## Acknowledgements

This work was supported by the V Foundation for Cancer Research (V2020-010, to A.C.P.), and NIGMS grants T32-GM135095 (S.C.P.) and K12-GM000678 (A.E.P.). We thank Clemens Grassburger and David Weinstock for helpful discussions, and David Weinstock for cell cultures.

## Methods

### Cell culture

PTCL cell lines were a gift from Dr. David M. Weinstock. Cell line identities was verified through short tandem repeat (STR) profiling by LabCorp and Cellosaurus CLASTR 1.4.4 database (**Table 1**). Cell lines were routinely tested for Mycoplasma and were negative (Lonza MycoAlert). Cells were cultured by standard methods in RPMI 1640 (Gibco) supplemented with Fetal Bovine Serum (Sigma) and Penicillin-Streptomycin (Gibco), at 37 °C and 5% CO_2_ (**Table 1**). Live cell densities were counted by BioRad TC20 and trypan blue.

**Table 1:** Growing conditions for 7 T-cell lymphoma cell lines. ALCL = anaplastic large cell lymphoma, ALK = anaplastic lymphoma kinase, FBS = fetal bovine serum, IL-2 = interleukin-2, NK = natural killer, NOS = not otherwise specified, STR = short tandem repeat, T-LGL = T-cell large granular lymphocytic leukemia. * STR profile not in Cellosaurus CLASTR 1.4.4 database

| Cell Line | Subtype | STR Profile Match | Supplements | Seeding Density (cells / mL) | Treatment naïve? |
| --- | --- | --- | --- | --- | --- |
| DL-40 | ALK-neg ALCL | 100% | 20% FBS | 10<sup>5</sup>; split every 3 days | Yes |
| KHYG-1 | NK | 100% | 10% FBS + 100U/mL IL-2 | 10<sup>5</sup>; split every 3 days | Yes |
| Ki-JK | ALK+ ALCL | 100% | 20% FBS | 2×10<sup>5</sup>; split every 2 days | Yes |
| MOT-N1 | T-LGL | 96% | 20% FBS | 10<sup>5</sup>; split every 3 days | Yes |
| MTA | NK | 97% | 10% FBS | 10<sup>5</sup>; split every 3 days | Yes |
| SMZ-1 | PTCL-NOS | N/A* | 10% FBS | 10<sup>5</sup>; split every 3 days | No<sup>38</sup> |
| SU-DHL-1 | ALK+ ALCL | 100% | 10% FBS | 10<sup>5</sup>; split every 3 days | Yes |

### Chemotherapy Preparation

4-hydroperoxycyclophosphamide (4-HC) was purchased from Niomech GmbH (D-18864) and all other chemotherapies were purchased from MedChemExpress. Prodrugs prednisone and cyclophosphamide were substituted with active species (prednisolone in place of prednisone and 4-HC in place of cyclophosphamide).

### Measurement of drug interactions by the Bliss model

Protocols for dose response measurements were as previously described^9^ and detailed below.

#### Cell plating

Cells at a density of 133,333 cells/mL were plated into 384-well black-bottomed plates (Nunc #164564) at 30 μL/well (corresponding to 4,000 cells/well), using a Thermo Fisher Multidrop Combi. After drug treatment, plates were incubated at 37 °C and 5% CO_2_ (Heracell VIOS 160i) with secondary containment in plastic tubs lined with sterile wet gauze to minimize evaporation. Two separate wells in a 24-well plate were plated with 1 mL cells each at the same density, to monitor growth rate over the course of the experiment. Live cell density in one well was counted at the time of drug administration, and cell density in the second well was counted at the end of the experiment to calculate growth rate in the absence of therapy.

#### Drug administration

Drugs were administered by a Tecan D300e digital drug dispenser. Wells were randomized during drug administration and re-organized during data analysis to avoid systematic spatial bias on the plates. Drugs were administered as single agents, pairwise, triplicate, and quadruplicate combinations in concentration ratios determined by their C_sustained_ values. C_sustained_ refers to a clinically relevant concentration based on measurements in patients’ serum up to 6 hours after administration^12^; these are: 4-HC, 15 μM^9,39^; doxorubicin, 150 nM^9^; prednisolone, 5 μM^40,41^; vincristine 5 nM^9^. Dose response measurements spanned concentrations from 0% to 500% C_sustained_ in log-spaced steps. Cells were incubated with drugs for 72 hours, which spans the *in vivo* elimination half-lives of these drugs^42^.

#### Measuring relative viability

After drug incubation, cell viability was quantified with Promega CellTiter-Glo (1:1 dilution in PBS) at 25 μL/well to visualize ATP levels by luminescence. CellTiter-Glo was administered with Multidrop Combi and plates were incubated for ten minutes. Plates were centrifuged at 200g for 5 minutes to eliminate air pockets. Luminescence was measured by BMG CLARIOstar reader using an aperture to reduce well cross-talk to below 10^−4^. Serial dilution of live cells confirmed a linear dynamic range over 4 orders of magnitude (100% to 0.01% relative live cell count) (**Supplemental Figure 1**).

#### Replicates

Measurements consisted of four technical replicate wells per drug concentration per plate, with two biological replicates comprising independently propagated cultures, for a total n=8 per data point.

#### Analysis

Relative viability expected by the Bliss Independence model was calculated by multiplying the relative viabilities produced by each single drug in a combination.

### Measurement of drug interactions by isobologram analysis

Cell plating, drug treatment, and viability measurements were performed as above.

#### Drug concentrations

For isobologram analysis, two-dimensional gradients of drug concentrations were prepared across a 11X11 grid of wells. Concentrations decreased in log-spaced steps across a 100-fold range, chosen to span a range from negligible inhibition to strong killing. Control wells with no treatment and single-agent dose responses were included on each plate.

#### Replicates

Two technical replicates were repeated in each of two biological replicates for a cumulative n=4. The precision of automated liquid-handling produced high consistency between biological replicates (**Supplemental Figure 1**).

#### Analysis

Relative viability at each concentration was an average of 4 independent experiments. A nearest-neighbor median filter was applied to relative viability across two-dimensional dose response surfaces. Isobolograms are plotted with contours highlighting 50%, 20%, and 5% relative viability.

### MuSyC Analysis

To overcome the limitations of single drug interaction metrics, we also applied a model that synthesizes Bliss’ and Loewe’s models, titled Multidimensional Synergy of Combinations (MuSyC)^17^. MuSyC distinguishes between drug response curves changing in potency (A) versus efficacy (B). To apply MuSyC to combinations of 3 or more drugs, we adapted the framework to quantify changes in dose response compared with that predicted by the Bliss model (**Supplemental Figure 2**). In this procedure, we applied shifts in potency (A) and efficacy (B) to the Bliss predicted response, to identify which parameters produce the best fit to the experimentally observed response, thereby quantifying how drug interactions change potency, efficacy, or both. If multiple values of A and B could produce similar best fits, we chose those with a consistent direction of change (e.g. increased potency and increased efficacy). This was necessary to exclude implausible claims of opposite interactions that cancel out and have no effect (e.g. increased potency and decreased efficacy).

### Sequential versus concurrent CHOP treatments

MTA cells were plated at 500,000 cells/mL in 10mL in 25cm^2^ flasks (Nunc EasYFlasks). Three independent cultures were propagated for each of five dosing conditions. In the control condition, two cycles of CHOP were administered on Days 0 and 12 at equipotent concentrations (C: 1.2 μM, H: 10 nM, O: 0.4 nM, P: 0.4 μM). Equipotent concentration ratios were defined by the monotherapy concentrations required for 12 days of growth suppression. Monotherapy concentrations were each lowered to 40% when constituting an equipotent 4-drug combination (full doses of each monotherapy would sterilize the flask), which corresponded to ≈10% C_sustained_. Prednisolone monotherapy is not inhibitory in PTCL cell lines but was administered at matching C_sustained_. For sequential treatment conditions, a three-drug cocktail was administered on Days 0 and 12, and the ‘offset’ drug was administered as a single agent on Days 6 and 18, at the same concentrations as the CHOP control. After every three-day treatment period, cells were washed twice by centrifugation, removal of supernatant, and resuspension in 10mL of drug-free media. Live cell count was measured (BioRad TC20) and the cells were transferred to a new 25 cm^2^ flask. Average live cell count was calculated from the three flasks per condition (n=3).

### Sequential versus concurrent CH treatments

Experiments were performed as described above with one cycle of therapy at concentrations of C: 2μM, H: 20 nM, and C: 1μM, H: 10 nM for the ‘half-dose’ condition.

### Adaptive Resistance

MTA cells were plated at 200,000 cells/mL in 50 mL for each of four 175 cm^2^ flasks. One control flask received no treatment, and others were treated with single agent 4-HC (2 μM), doxorubicin (20 nM), or vincristine (0.35 nM). After three days of treatment, cells were washed twice, resuspended in drug-free media, and live cell count was measured (as described above). Cells recovered in drug-free media for 4 days, plated at 100,000 cells/mL in 50mL in 175 cm^2^ flasks. On Day 7 post-treatment, cells were plated in 384 well plates (one plate per pre-treatment condition) and the dose response assays were performed as described in ‘Measurement of drug interactions by the Bliss model.’

### Model of ultrasensitive response and heterogeneity

Model parameters are in **Table 2**. The ‘Linear-Quadratic’ response equation (**Eqn 1**) is used in radiation oncology to describe the relationship between cell survival (*S*) and dose (*D*). The ‘linear’ component of dose response (α *D*) corresponds to the probability that cells die from a single DNA damage event. The ‘quadratic’ component (β *D*^2^) corresponds to the probability that cells die from an accumulation of DNA damage. A high ratio β/α produces a more ultrasensitive response, which favors ‘hypofractionated’ treatment schedules. Here, we apply the LQ model to cytotoxic chemotherapies where treatment ‘fractionation’ is also relevant^37^.

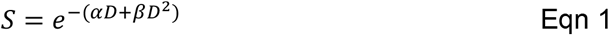

**Table 2:** Parameters for the ultrasensitive drug response model.

| Parameter | Value | Definition | Source |
| --- | --- | --- | --- |
| $\tau_{\text{death}}$ | 2.5 | Half-life of dying cells | Measured (**Supp. Fig. 4a**) |
| $\tau_{\text{growth}}$ | 2.5 | Doubling time of growing cells | Measured (**Supp. Fig. 4b**) |
| $i$ | -0.8 | CH drug interaction – less than additive | Measured (**Supp. Fig. 4f**) |
| $\mu$ | 0 | Sensitivity mean | Default |
| $\sigma$ | 0.42 | Sensitivity standard deviation | Fit to experimental data<br>(**Supp. Fig. 4g**) |
| $\beta$ | 0.4 | Quadratic component LQ model | Fit to experimental data<br>(**Supp. Fig. 4g**) |
| $\alpha$ | 0.1 | Linear component LQ model | Default |
| $D_C$ | 2.6 | Dose of 4-H-cyclophosphamide | Measured (**Supp. Fig. 4d**) |
| $D_H$ | 1.8 | Dose of doxorubicin | Measured (**Supp. Fig. 4d**) |
| $P_0$ | $5 \times 10^5$ | Initial number of tumor cells | Used in experiments |

A linear dose response shown in **Figures 5 and 6b** is:

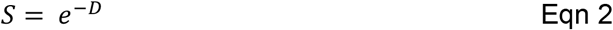

A ‘diminishing returns’ dose response (sub-linear) shown in **Figure 6b** is:

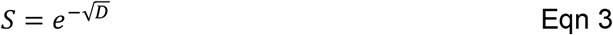

To model cellular heterogeneity in drug sensitivity, we describe a population of tumor cells with log-normally distributed drug sensitivity. That is, ‘x’ is normally distributed (μ = 0, σ = 0.42) and 10^x^ is the drug sensitivity parameter, which is multiplied by the dose response (α *D* + β *D*^2^) to describe response in cells having greater or lesser drug sensitivity (**Eqn 4**).

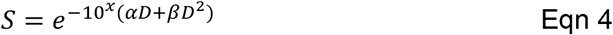

Average survival of a cell population with *N* subpopulations having survival is 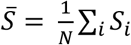. **Figure 4d** shows average survival when quantified as ‘log-kills’, defined as λ = −log_10_(S). The average number of log-kills is therefore

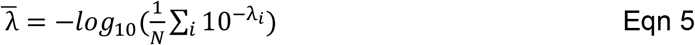

To calculate the fraction of cells that survive drug treatment, for a population of cells with heterogeneous drug sensitivity, the dose response equation is integrated over the drug sensitivity distribution (**Eqn 6**).

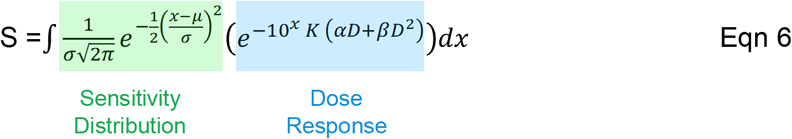

When modeling response to multiple drugs (e.g. A and B) with drug interactions, the dose term *D* is replaced by an effective combined dose *d* (**Eqn 7**) which may be greater or lesser than *D*_A_ + *D*_B_ depending on the sign of a drug interaction term *i*:

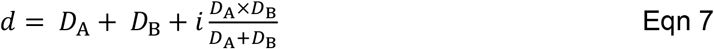

For any value *i*, **Eqn 7** yields a consistent ‘Combination Index’ (CI) at all magnitudes of effect, with *i* =0 being additive, *i* >0 being synergy (higher effective dose), and *i* <0 being antagonism (lower effective dose). This is proven below for the case of an equipotent combination (*D*_A_ = *D*_B_):

Effective combined dose × Combination Index = Sum of doses

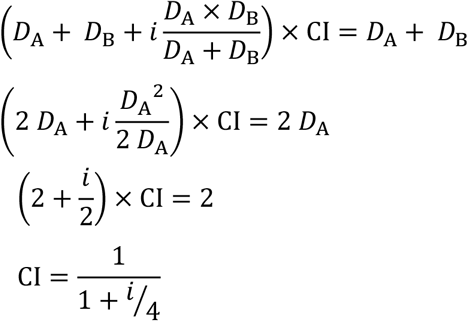

To model sequential treatments, it is not sufficient to multiply individually calculated values of S, as this would not track the consequence of the first cycle killing more drug sensitive cells. Instead, sequential dose response functions are multiplied within the integral:

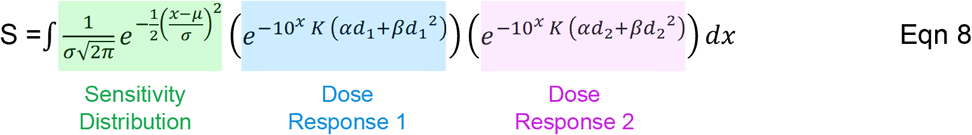

To model response kinetics, cells that survive drug treatment grow exponentially with a doubling time τ_growth_ (experimentally observed), named the growing population G(t) (**Eqn 9**). The population of cells dying from chemotherapy, H(t), are modelled as dying exponentially rather than instantly, because experimental measurements at sterilizing doses of chemotherapy reveal exponential death kinetics, with half-life τ_death_ (experimentally observed) (**Eqn 10**). Experimental cell counts measure both live, growing cells and also live cells that are in the process of dying. Therefore, the final model output, total live cell population, is the sum of growing and dying cell populations (**Eqn 11**).

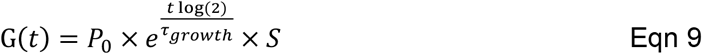

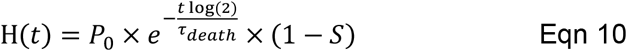

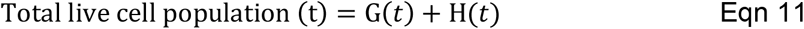

Importantly, the model accounts for the change in sensitivity across the cell population caused by drug selection. As a consequence of repetitive treatments, the drug sensitive populations are the first to die, and as a result, the total cell population grows progressively less drug sensitive. The model accounts for this phenomenon by quantifying the additional cell kill fraction from the previous treatment instead of assuming each treatment has an equivalent cell kill fraction.

When hypothetical drug pair ‘A+B’ was modeled in **Figure 6a**, each single dose was modeled at D = 3. Similarly, in **Figure 6b**, the hypothetical single agent was modeled at D = 12 for each single dose.

