## Supplemental Figures 1 to 6 for "Ultrasensitive response explains the benefit of combination chemotherapy despite drug antagonism"

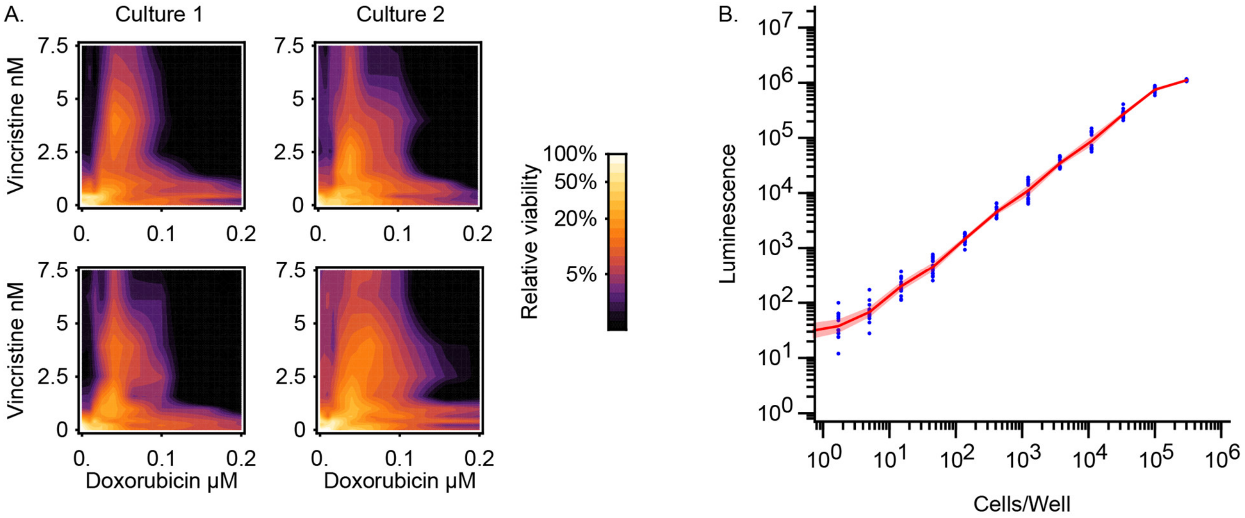


**Supplemental Figure 1. Reproducibility and dynamic range of *in vitro* drug interaction experiments.** A) Consistency of measured isobolograms among biological replicates (independently propagated cultures; left and right columns) and among technical replicates (different microtiter plates; top and bottom rows). Cell line = MTA, n=1 per plot here, with each of 4 plots averaged to produce isobolograms in **Figure 1**. B) Dynamic range of CLARIOstar microplate reader is linear over 5 orders of magnitude in cell density ($10^1$ to $10^5$ Ki-JK cells per well). Red shaded region represents 95% confidence interval of population mean (n=16).

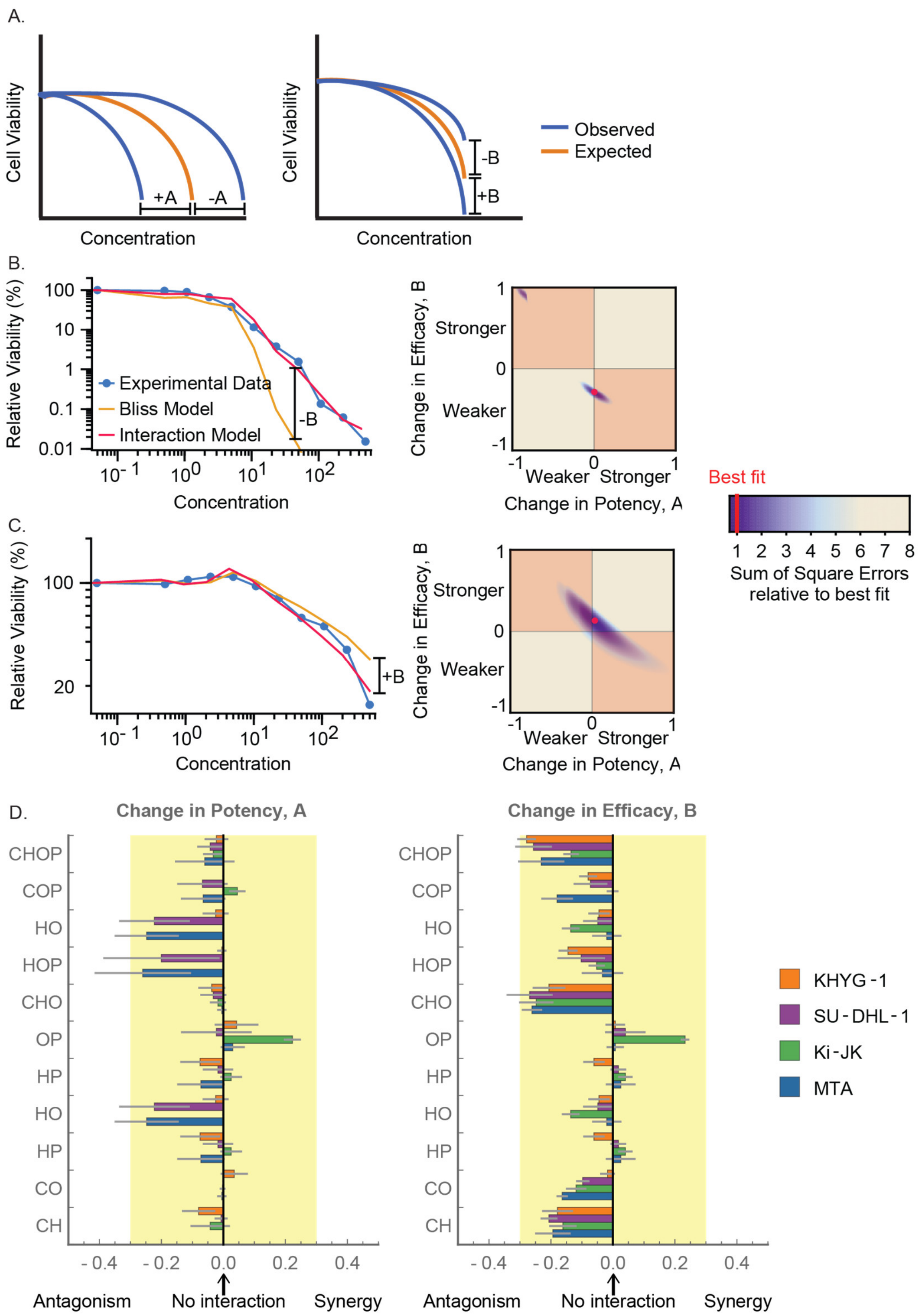


**Supplemental Figure 2. Drug interactions in CHOP using MuSyC method to distinguish potency and efficacy shifts.** A) The MuSyC method analyzes drug combination effects to distinguish between shifts in potency and shifts in efficacy. Shifts in potency from the Bliss model (orange) to observed data (blue) are represented by parameter A, where positive values indicate superior potency and negative values indicate inferior potency. Changes in efficacy are represented by parameter B, where positive values indicate greater inhibition, and negative values indicate less inhibition. B) Example of method for a drug combination with lower efficacy than predicted by the Bliss model. The interaction model (red) is fit from the Bliss model to the experimental data across the length of the curve. The quadrant plot shows the A and B parameters that confer the best fit determined by sum of squares errors. Stronger fits are colored darker blue and the best fit is marked in red. Data shows drug combination CHOP in SU-DHL-1 T-Cell lymphoma cell line, n=8. C) Example of method for a drug combination with higher efficacy than Bliss model predicts. Data shows drug combination leuocorovin-fluorouracil-irinotecan in Lovo Colorectal cancer cell line, n=8. D) Shifts in potency and efficacy for all 11 combinations of agents in CHOP, for each of four Peripheral T-Cell Lymphoma cell lines: KHYG-1, SU-DHL-1, Ki-JK, MTA. Yellow region marks a two-fold difference in potency or efficacy compared with the ‘no interaction’ null hypothesis. Error bars are 95% confidence intervals, n=8.

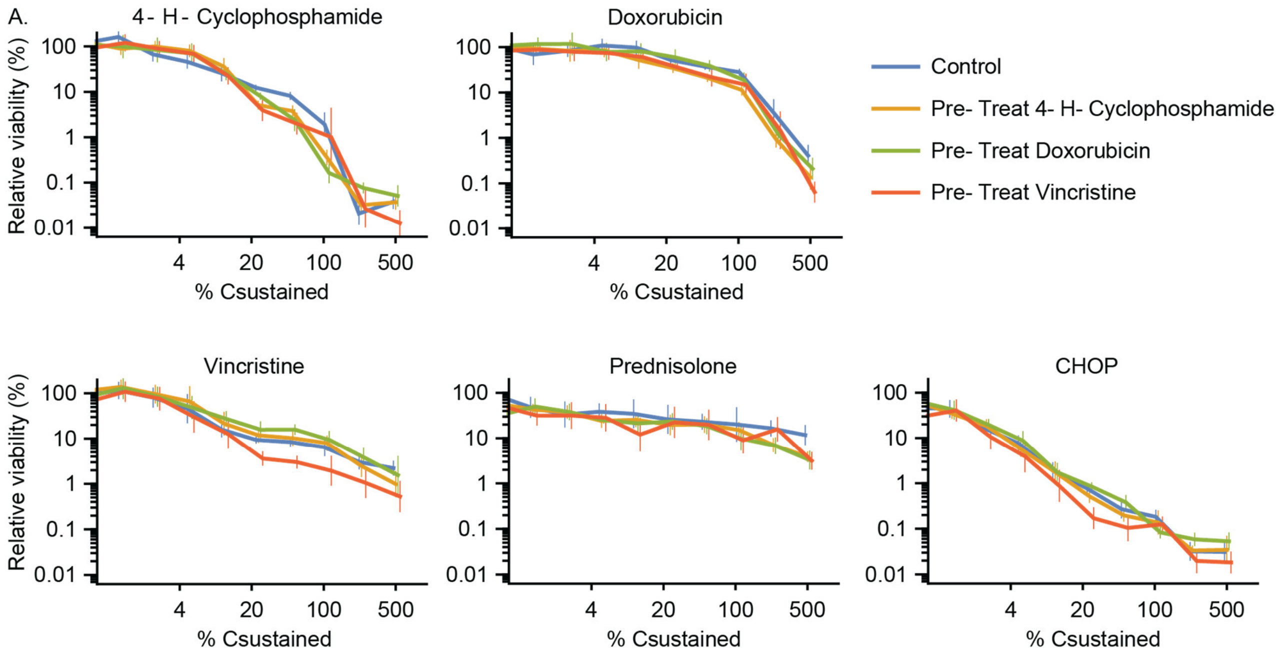


**Supplemental Figure 3. Short-term adaptive resistance does not affect sequential treatments.** MTA cell cultures were treated for 3 days with either 4-H-cyclophosphamide, doxorubicin, vincristine, or no treatment (control). Drugs were administered at concentrations producing 50% growth inhibition. Following treatment, cultures were washed of drug and recovered in drug-free media for 4 days. After recovery, dose responses were measured for 3-day treatments with 4-H-cyclophosphamide, doxorubicin, vincristine, prednisolone, and the CHOP combination. Pre-treated cultures were not more resistant to chemotherapy than control cultures. Error bars represent 95% confidence intervals, n=8.

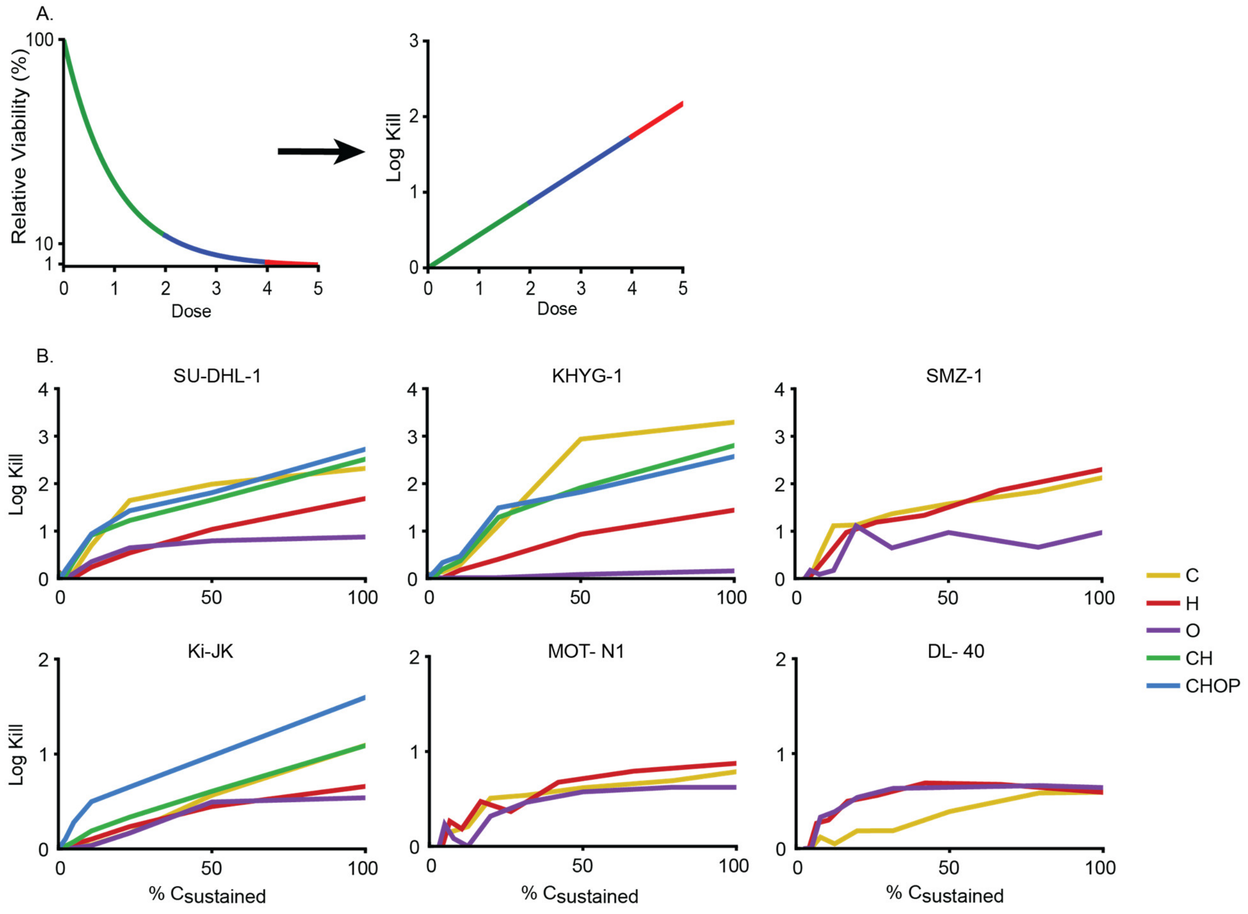


**Supplemental Figure 4. Experimental population dose responses on log-kill scale.** A) Many chemotherapies exhibit exponential dose responses in hematologic cancers (left), which appear linear when plotted on a logarithmic scale (right). The first 90% cytotoxic effect is shown in green (reaching 1 log-kill); the next 90% cytotoxic effect is shown in blue (reaching 2 log-kills); the next 90% cytotoxic effect is shown in red (reaching 3 log-kills). B) Log-kills produced by CHOP and its components at concentrations from 0 to 100% $C_{sustained}$. n=8 for SU-DHL-1, KHYG-1, and Ki-JK; n=4 for SMZ-1, MOT-N1, and DL-40.

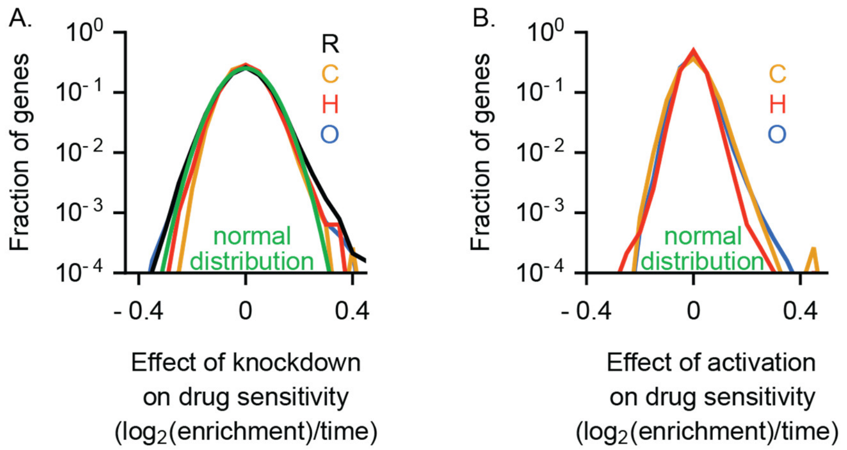


**Supplemental Figure 5. Previously published CRISPR inhibition (A) and CRISPR activation (B) data shows log-normally distributed drug sensitivities across a heterogeneous tumor cell population.** When heritable heterogeneity was induced via CRISPR inhibition or activation, cell populations had log-normally distributed sensitivities. A) Genome-wide CRISPR inhibition screen was performed in dCas9-KRAB expressing Pfeiffer cells (diffuse large B-cell lymphoma) treated with single agent rituximab (R), 4-H-cyclophosphamide (C), doxorubicin (H), and vincristine (O). B) Genome-wide CRISPR activation screen was performed in dCas9-VP64 expressing K562 cells (chronic myeloid leukemia) and treated with single agent 4-H-cyclophosphamide (C), doxorubicin (H), and vincristine (O). Data publicly available from Palmer et al. eLife (2019)[9].

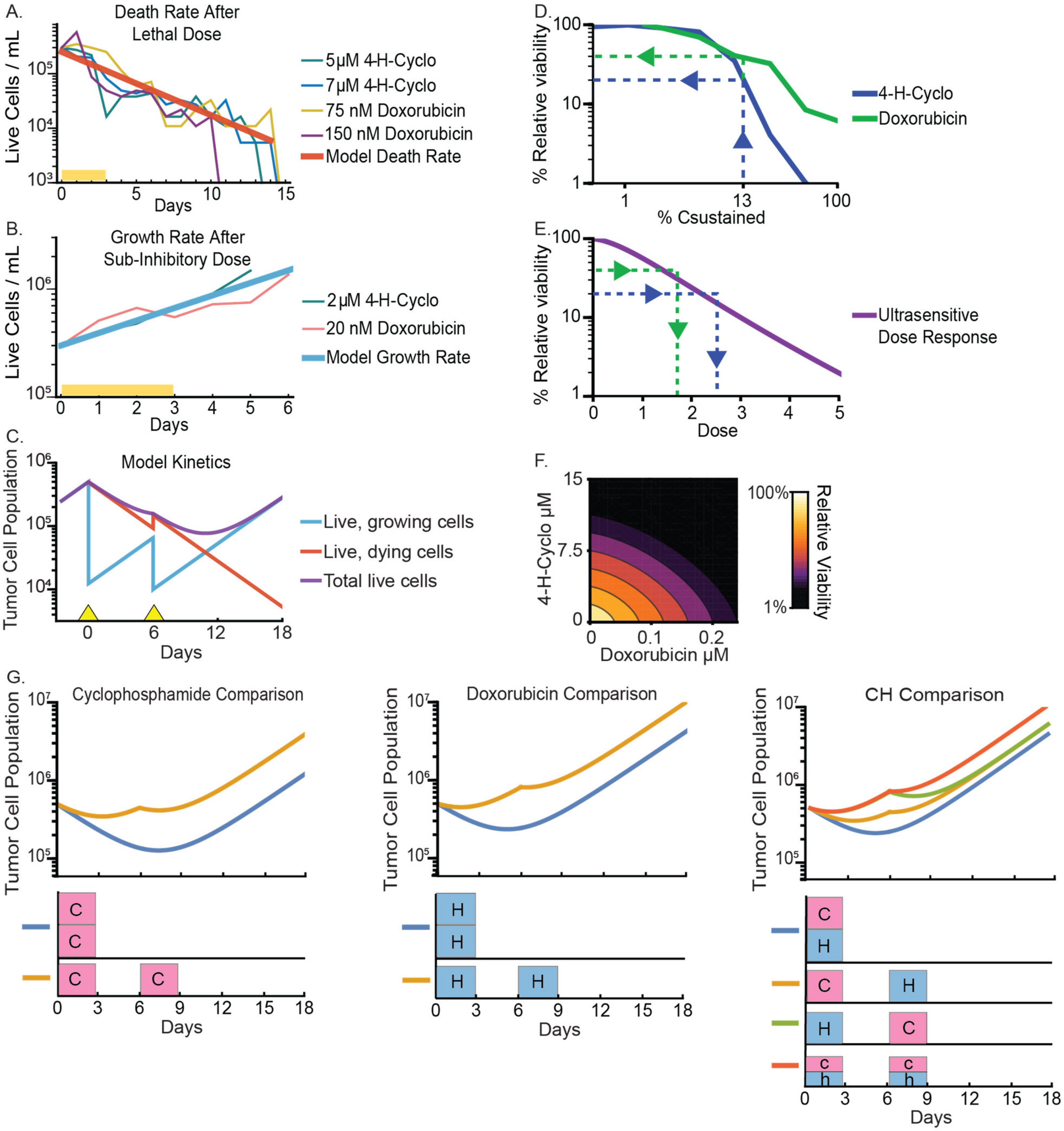


**Supplemental Figure 6. Model parameters informed from experimental data.** A) The model's death rate was determined by treating MTA cells with lethal doses of 4-H-cyclophophamide and doxorubicin and observing that the cell population halves every 2.5 days, n=1 per condition. Cells were incubated with drugs for 3 days (represented by yellow bar) before being washed off and replaced with drug-free media. B) The model's treated growth rate was determined by treating MTA cells with the same drug concentrations used in the concurrent vs sequential dosing experiments and observing that cells double every 2.5 days, n=1 per condition. Cells were incubated with drugs for 3 days (represented by yellow bar) before being washed off and replaced with drug-free media. C) At each simulated drug treatment (marked with yellow triangles), the model calculates the fraction of cells that will survive and the fraction that will die. Each respective cell population is modelled as growing or dying at the experimentally determined rates. Total live cell count is calculated by adding the number of growing cells with the number of dying cells over time. D) 4-H-cyclophosphamide yields about 20% viability and doxorubicin yields about 40% viability at 13% $C_{sustained}$, the concentration of drug that was used in the concurrent vs sequential administration experiments (2 μM and 20 nM respectively), n=8 per condition. E) To convert experimental concentrations to dose values for the model, the corresponding dose values at 20% and 40% viability were estimated to be 1.8 for doxorubicin and 2.6 for 4-H-cyclophosphamide. F) The CH isobologram created by the model matches the experimentally determined isobologram for MTA cells in **Figure 1b** when the drug interaction term is set to a sub-additive value of i = -0.8. G) Model outputs of tumor cell populations over time were fit to the experimental data from **Figure 3c+d** to fit the level of ultrasensitivity (β = 0.4) and the level of heterogeneity in drug sensitivity (σ = 0.42). The simulated dosing schemes are visualized under each dose response plot with 4-H-cyclophophamide in pink and doxorubicin in blue.
